# Protein structures as shapes: Analysing protein structure variation using geometric morphometrics

**DOI:** 10.1101/219030

**Authors:** Jose Sergio Hleap, Christian Blouin

## Abstract

A phenotype is defined as an organism’s physical traits. In the macroscopic world, an animal’s shape is a phenotype. Geometric morphometrics (GM) can be used to analyze its shape. Let’s pose protein structures as microscopic three dimensional shapes, and apply principles of GM to the analysis of macromolecules. In this paper we introduce a way to 1) abstract a structure as a shape; 2) align the shapes; and 3) perform statistical analysis to establish patterns of variation in the datasets. We show that general procrustes superimposition (GPS) can be replaced by multiple structure alignment without changing the outcome of the test. We also show that estimating the deformation of the shape (structure) can be informative to analyze relative residue variations. Finally, we show an application of GM for two protein structure datasets: 1) in the *α*-amylase dataset we demonstrate the relationship between structure, function, and how the dependency of chloride has an important effect on the structure; and 2) in the Niemann-Pick disease, type C1 (NPC1) protein’s molecular dynamic simulation dataset, we introduce a simple way to analyze the trajectory of the simulation by means of protein structure variation.

## Introduction

Geometric morphometrics is a collection of approaches for the multivariate statistical analysis of Cartesian coordinate data [Slice, 2007]. The “geometry” referred to by the word “geometric” is the estimation of mean shapes and the description of sample variation of shape using the Procrustes distance (Kendall’s shape space) [Rohlf, 2002]. It is mainly based on landmarks which are “*discrete anatomic loci that can be recognized as the same loci in all specimens of study*” [Zelditch et al., 2004, p.443], and must: be homologous anatomical loci, not alter their topology or position relative to other landmarks, provide adequate coverage of the morphology, be consistently assigned, and lie within the same plane [Zelditch et al., 2004].

Landmark data is more informative than traditional data (linear measurements in the geometric morphometrics context) since its coordinates also contain positional information and thus geometric structure. Once homologous landmarks are assigned, “noisy” factors affecting the dimensionality and degrees of freedom of the possible shape analysis are removed by means of generalized Procrustes superimposition (GPS). Such factors being rotation, translation and size, and are dealt with by [Adams et al., 2004, Zelditch et al., 2004]:

1. Assign homologous landmarks to meaningful and descriptive parts of the shape.
2. Center each configuration of landmarks at the origin by subtracting the coordinates of its centroid from the corresponding (X or Y) coordinates of each landmark: Removing positional variation by translating each centroid to the origin.
3. Scale the landmark configuration to unit centroid size by dividing each coordinate of each landmark by the centroid size of the configuration.
4. Set one configuration as a reference and rotate the other configurations to minimize the summed squared distances between homolog landmarks, thus removing rotational variation.

The above method can be expressed as [Rohlf and Slice, 1990]:

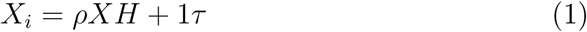

where matrix *X* is the original configuration; *ρ* is the scaling done to *X*; 1*τ* is the translation performed to *X*_*i*_ to a reference position; and *H* is a rotation (with an angle of rotation *θ*) matrix of the form:

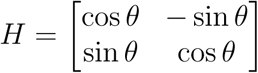

Then, the set of all matrices representing the landmark configurations (configuration space), becomes the shape space, and its dimensions are given by:

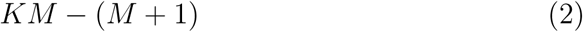

where *K* is the number of landmarks, and *M* the number of dimensions in each landmark. After removing the effects of size, rotation and translation, 2*K −* 4 degrees of freedom are left for 2D data and 3*K −* 7 for 3D data [Zelditch et al., 2004].

The comparison and analysis of the superimposition is based in the the Procrustes distance (*D*_*P*_). GPA applies the Procrustes analysis method to align a population of shapes instead of only two shape instances [Dryden and Mardia, 1998]. The Procrustes distance is the square root of the sum of squared differences between the positions of the landmarks in two optimally superimposed configurations (*C*_1_ and *C*_2_) [Rohlf, 2002]:

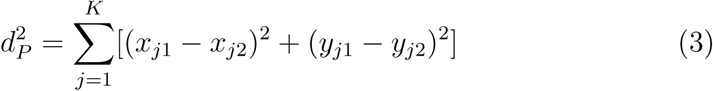

where 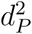 is the squared Procrustes distance, and *K* is the number of landmarks. This metric can then be used in several statistical multivariate analyses to attain differences in shapes, clustering, changes in time, test symmetry, etc. Here shapes can be treated as a single point in a multidimensional space, and therefore the information can be summarized in an efficient way using standard multivariate techniques. Some traits can also be treated independently in the analyses, extracting information of particular aspects of the shape.

This paper shows how these geometric morphometric methods and applications can be use to analyze protein structure variation. It will start by proposing a method to abstract a protein structure as a shape. Then a demonstration that the scaling applied in the General Procrustes Superimposition (GPS), which does not apply to protein structures, can be omitted without changing the results will be shown. It will also be shown that GPS can be replaced by a Multiple Structural Alignment (MStA), which handles the rotation and translation part of the transformation. A simple method to analyze individual residue variation within this context and relate it to structural issues will be introduced. Finally, we will show general applications of GM-like methods for protein structure variation analysis in two datasets of actual protein structures.

## Material and methods

### Abstracting a protein structure as a shape

A landmark will be defined as the centroid of each homologous residue, and is defined by (x,y,z):

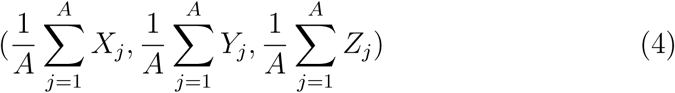

where *A* will be the number of heavy atoms that constitutes the side chain of a residue including the *C*_*α*_. This procedure takes into account only homologous residues assigned by a structural alignment using the MStA application MATT [Menke et al., 2008]. Only ungapped homologous sites will be taken into account.

### Testing the effect of scaling in protein datasets

The HOMSTRAD (386 datasets) and SABmark (425 datasets) superfamily subsets reported in MATT’s paper [Menke et al., 2008] were used. The former database was design to store structures based on the quality of the X-ray analysis and accuracy of the structure, while the latter database was designed to test multiple alignment problems. Residue homology was determined by MATT-reported alignment [Menke et al., 2008, http://groups.csail.mit.edu/cb/matt/]. The centroid’s coordinates (see section Abstracting a protein structure as a shape) for each homologous residue were computed and stored. To the resulting centroid’s coordinates file, GPS with and without scaling were performed with the R [R, 2011] package “Shapes” [Dryden, 2011]. A correlation analysis was performed graphically and using the Pearson test for correlation using the MatPlotLib [Hunter, 2007], and Scipy [Jones et al., 2001] libraries in Python.

### Testing GPS versus structural alignments in protein datasets

As in section Testing the effect of scaling in protein datasets, the HOMSTRAD and SABmark datasets where used to test the effect of aligning the protein structures with GPS or the flexible MStA implementation in MATT. The matrix of pairwise RMSD per dataset were computed, and a correlation analysis was made following the methods of section Testing the effect of scaling in protein datasets.

### Analysis of the deformation in superimposed structures/shapes

The inter-landmark distance matrix (form configuration) is computed using the Euclidean distance for each entry in *m* dimensions:

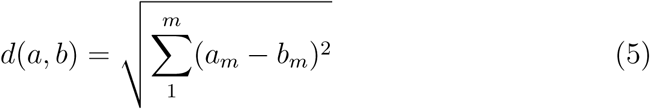

where *d*(*a, b*) stands for the Euclidean distance between variables *a* and *b*. Therefore the form matrix (*F M*) is [Claude, 2008]:

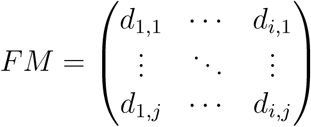

*F M* is then a square symmetric matrix, with zeros in the diagonal entries. If two forms (shapes in the inter-landmark framework) are identical, they will have the same entries in the *F M* matrix. The matrix of differences in form [The form difference matrix or FDM as named by Claude, 2008] between two configurations *S*1 and *S*2 is given by:

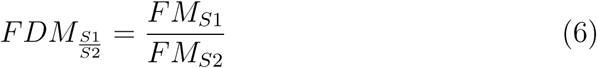

The score of the most influential point (*I*) in the data can be computed by adding the sum of the differences to the median value per column (variable) and ranking the positions. This can be computed as:

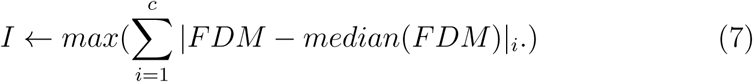

where *c* is the column index, and FDM is the form difference matrix.

However, as shown in equation 6, this FDM is the representation of the difference between two shapes. We can generalize by summing the residuals of all shapes versus a hypothetical mean shape. For simplicity this can be calculated as the per-variable per-dimension average. In other words, the average of each dimension of each landmark. This approach will then return a Form Difference (FD) value per landmark, however; this value is not bounded and is difficult to interpret. For this reason we scaled the resulting FD vector 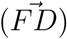 such that it is bounded from −1 (least variation) to 1 (highest variation) with:

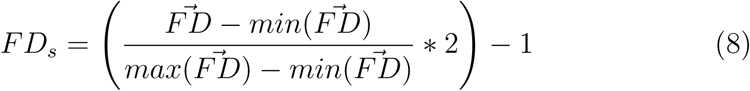

To illustrate how this works, a simulation of 500 hexagons was performed. Giving an initial shape, for each point and each dimension in the point, a distribution of random normal numbers is created. This distribution has a given standard deviation (0.05 for regular points, 0.005 for low variation points, and 0.2 for highly variable points). The mean was set to 0. This distribution per point and per dimension was then added to the original shape, creating the simulated dataset with controlled variation. To explore a more complex system than the hexagon, a protein simulation was also performed in the same fashion as for the 2D case. In this case more extreme points were used for visualization purposes, being 0.0005 the lowest variation and 0.5 the highest. The protein used for the simulation and visualization was the Porcine Pancreatic Amylase (PPA) with pdb code 1PPI.

To test the variation in an evolutionary perspective, a protein structure sampling was performed. A PFAM [Finn et al., 2010] seed alignment of the *α*-Amylase family was gathered and used to seed a PSI-BLAST [Altschul et al., 1997] search. The PSI-BLAST search was restricted to structures available at the protein data bank (http://www.rcsb.org/pdb/). There were 135 structures gathered in total (Table S1 in Supplementary data) whose homology and membership to the *α*-amylase family (the Glycoside Hydrolase Family 13, GH13) was guaranteed. After the sampling, the methods mentioned above were applied to the dataset and analyzed with *F D* and *F D*_*s*_.

### Statistical analysis of protein structural data

A multidimensional scaling (principal coordinates) analysis and *F D*_*s*_ estimation and analysis were performed to two real datasets. The first one, as described above, is a dataset of 135 structures of the *α*-Amylase family. With this dataset, the variation observed is assumed to be driven by the evolutionary history of the structure.

The second dataset is a molecular dynamic simulation of the Niemann-Pick, type C1 N-terminal domain (NPC1), in solution. The software GROMACS 4.5 [Hess et al., 2008] was used with the force fields OPLS-AA/L [Jorgensen and Tirado-Rives, 1988] for the protein, and the TIP3P [Jorgensen et al., 1983] for the water molecules. The data was collected every 20 picoseconds for 100 nanoseconds discarding the first 10 nanoseconds of simulation to achieve stability. This process was performed using a 24-core GPU-enabled workstation. Each sample was treated as an individual observation for the subsequent analysis, and the data are extracted and processed as explained in sections above. In this dataset two simulations were performed: with Cholesterol (NPC1’s ligand) bound to the structure, and another one without the ligand.

## Results and Discussion

A protein fold can be essentially defined as a 3D geometric shape. Sequence analyses help to understand some trends, but explain little about geometry. GM can be used to perform shape analysis from a geometric point of view. It also can be used to give insight into the phylogenetic relationships of the structures rather than the sequences. However, the application of GM to protein structures is not trivial. The scaling component of the Procrustes analysis has no conceptual equivalent for proteins. Since organisms grow, it makes sense to extract the size effect on shape in order to compare young with adults. On the other hand, in proteins the atoms do not stretch or grow, and therefore scaling [as proposed in Adams and Naylor, 2000, 2003] is not appropriate.

In [Adams and Naylor, 2000] and [Adams and Naylor, 2003] proposal, they:

- Abstract a residue as a landmark.
- Determine the homologous residues, using ClustalW [Thompson et al., 1994].
- Delete non-homologous sites.
- Perform morphometric analyses.

The use of sequence alignment without structural information to infer structural homology is not accurate since the amount of gaps that can be allowed in a loop region can be different than in other regions of the protein [Kann et al., 2005, Kjer et al., 2007], and therefore the definition of structural homology can be different as well. Moreover, since structures are more conserved than sequences, the alignment based on the structures allows a more reliable asssignment of homology in more distant clades [Wohlers et al., 2012].

In contrast, here we used protein structural alignment which has been worked on extensively [Kolodny et al., 2005, Hasegawa and Holm, 2009, Poleksic, 2011, Joseph et al., 2011, Shibberu et al., 2012]. In particular we used a flexible structure alignment method [MATT; Menke et al., 2008]. This approach strips out rotational and translational information as well as the variability induced by flexible hinges, which has been shown to be more accurate than rigid body superimpositions in the assignment of homology [Menke et al., 2008, Konc and Janežič, 2010, Nguyen et al., 2011, Daniluk and Lesyng, 2011, Joseph et al., 2012, Shah and Sahinidis, 2012, among others].

The abstraction of the residues and landmarks is similar to that in [Adams and Naylor, 2000] and [Adams and Naylor, 2003]; however, those papers did not fully describe the abstraction.

### The effect of scaling in GPS: Insights form HOMSTRAD and SABmark superfamily databases

In protein structure datasets it is expected that the scaling does not play a major role in the alignment of structures. To test such expectation, a correlation test between the two approaches was performed. The Homstrad dataset showed a correlation coefficient of 0.998, significant (*pval* < 0.001), and with an *R*^2^ of 0.997. The SABmark dataset also showed a significant and high correlation coeficient (*r* = 0.994*, pval* < 0.001) with an *R*^2^ of 0.987.

We have shown here not only that scaling is not conceptually acceptable in the context of protein structures (fixed lengths of atomic bonds), but that doing so won’t significantly affect the results, supported in high and significant correlation coefficients. From this point further, all GPS analyses made here will be referred as non-scaled GPS.

### Comparing MATT flexible alignment and GPS: Results from the HOMSTRAD and SABmark super family databases

To be able to compare between the MStA and GPS the two types of analyses have to be done independently but comparatively (e.g, using the same variables). However, GPS requires the assignment of homology of the landmarks. As mentioned in Methods, this homology is estimated by the multiple structural alignment. The GPS is an alignment itself, so providing a starting alignment might bias the result of the superimposition. To address this bias, the protein structures are aligned using MATT [Menke et al., 2008] for both approaches and the homologous residues’ index for each structure is recorded. For the GPS, the coordinates of the residues corresponding to the recorded indexes are used. This process is performed for each independent structure, therefore avoiding (or at least diminishing) the bias.

Since GPS aligns the structures based only on rotation and translation, it is logical to think of it as a rigid body superimposition. It does not allow any deformation of the structure (shape), like the flexibility that softwares like MATT have. Therefore, it is plausible to hypothesize that a flexible alignment will do better than GPS, as they do against non-flexible alignments [Menke et al., 2008, Konc and Janežič, 2010, Nguyen et al., 2011, Daniluk and Lesyng, 2011, Joseph et al., 2012, Shah and Sahinidis, 2012, among others]. Other flexible structural alignment softwares have shown a slightly better performance than MATT, however,the improvements are not significant [Joseph et al., 2012] and MATT returns more core residues than most of its competitors, as well as a statistical test of the “goodness” of the alignment [Menke et al., 2008].

In both databases (Homstrad and SABmark), the correlation coefficients were found to be greater than 0.99, showing that there is no difference in the use of either approach. This was an unexpected result since if a protein structure is allowed to bend, it is reasonable to expect a better fit. For the datasets explored this seems not to be the case. This observation can be explained if most of the analyzed datasets comprise single domain (which they do) proteins and are therefore more likely to behave as rigid bodies. Also, since GPS depends on the definition of homology inferred by an alignment provided by MATT, GPS could be functioning as a secondary alignment.

Because of this lack of difference, MATT alone is used in the rest of this manuscript.

### Form difference: Insights into residue variation

A sibling field to GM, *Dysmorphometrics* [Claes et al., 2012], can be used to explore the impact of outlier variables. Dysmorphometrics is in summary, “the modeling of morphological abnormalities” [Claes et al., 2012]. Such exploration can be performed by means of corrected maximum likelihood estimates approach [as in Claes et al., 2012] or by means of the Euclidean distance matrix analysis approach [Claude, 2008]. Here we use the latter since it is simpler and requires fewer parameters to be set.

Claude [2008], after the work of Lele and Richtsmeier [1992], proposed a way to examine the influence of landmarks in shape difference by calculating the sum of residuals from the median for each landmark given the *F D M* matrix. The landmarks that influence the most difference in shape would have a higher score which can be mapped to the given shape. In the context of geometric morphometrics, the FDM accounts for landmarks that have an excessive variation among a pair of shapes.

#### Shape simulation

To illustrate this effect, a hexagon simulation was performed where the variation among the observations is controlled. Figure 1 shows the plotting of 500 samples in this simulation.

**Figure 1:**
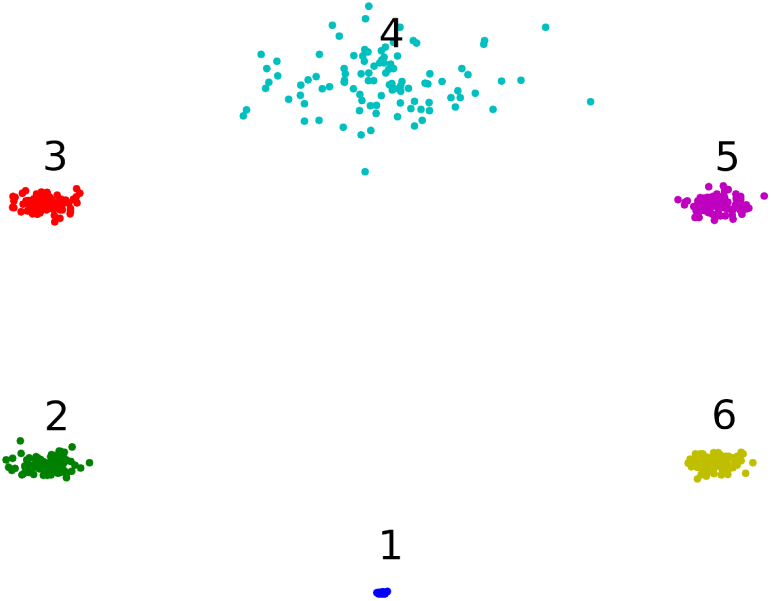
Hexagon simulation. Points 2, 3, 5, and 6 have a standard deviation of 0.05, while point 1 (low) have 0.005 and point 4 (high) have an standard deviation of 0.2. This plot was performed using the python library Matplotlib [Hunter, 2007].

Here a point with high variation (point 4 in Figure 1) and a point with minimal variation (point 1 in Figure 1) are introduced, along with four other points exhibiting an average variation in the context of geometric morphometrics [von Cramon-Taubadel et al., 2007].

Table 1 and Figure 1 show that the *F D*_*s*_ represents the overall influence of a point in the shape. Negative values of *F D*_*s*_ mean a more conserved point, being −1 the most conserved. On the other hand, positive *F D*_*s*_ values are related to more influential points, being +1 the most variable. From Table 1, one can also see that the “average” points are closer to the least variation than to the highest one, therefore displaying a negative tendency. At fist sight it might seem trivial to use the *F D*_*s*_ since the standard deviation (*sd*) seems to correlate with it. However, *F D*_*s*_ represents the influence of a landmark relative to the overall shape, as opposed to the *sd* which represents the variation at a single variable level. Variables with high *sd*s in a medium-*sd* neighbourhood will have very high *F D*_*s*_, while very low-*sd* points will have low *F D*_*s*_. This suggests that the *sd* can be used as proxy for *F D*. However, in a setting where all the points have high *sd*, the relation between *sd* and *F D* is not as direct. Moreover, for the relationship between *sd* and *F D*_*s*_ to be proportional, a model of isotropic variation in all dimensions is needed. Such model assumes an equal amount of variation at each landmark and at each dimension in each landmark. It also assumes that landmarks are independent, which is not a fair assumption in most shape analysis [Klingenberg, 2003].

**Table 1:** Scaled FD values for the simulation illustrated in Figure 1.

| Landmark | Standard deviation | Scaled FD |
| --- | --- | --- |
| 1 | 0.005 | -1 |
| 2 | 0.05 | -0.78 |
| 3 | 0.05 | -0.33 |
| 4 | 0.2 | 1 |
| 5 | 0.05 | -0.28 |
| 6 | 0.05 | -0.83 |

Non-scaled FD can be also used to screen a set of points in a shape to look for “Pinocchio” outliers, using statistical tests as the Dixon’s Q test or the Grubbs’ test for outliers.

#### *α*-Amylase simulation and evolutionary dataset

To test the form difference (FD) in more complex shapes, the *α*-Amylase dataset was used. First a simulation using the porcine pancreatic amylase (1PPI) was used to control the variation in two random residues (see section Analysis of the deformation in superimposed structures/shapes for details). Figure 2A shows the result of the FD analysis on the simulation.

**Figure 2:**
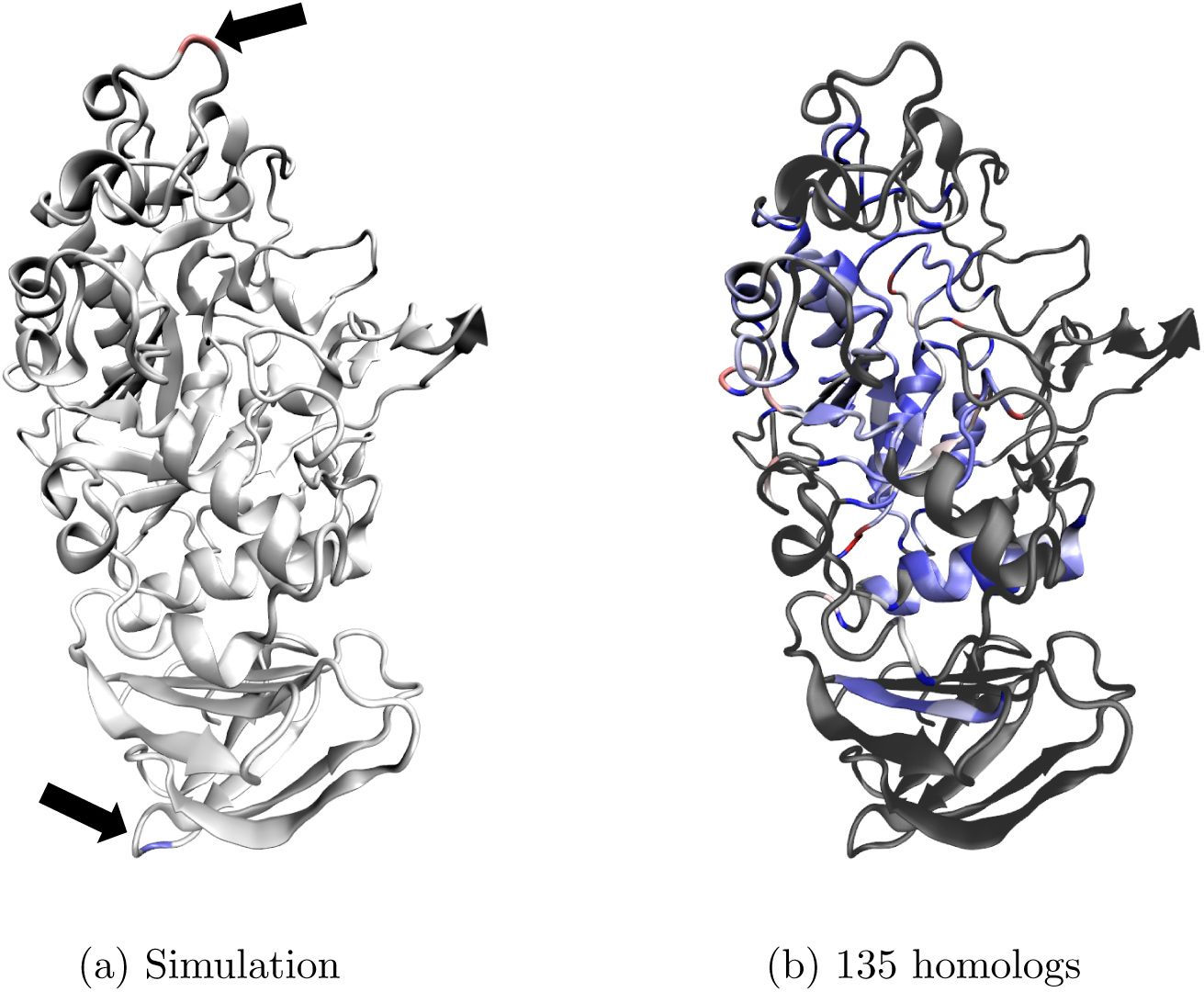
*F D*_*s*_ values mapped in the porcine pancreatic Amylase structure (PDB code: 1PPI). Red represents the highly variable, while blue the least variable. A) A simulation of the values (The locations of the points are selected at random and do not represent any biological meaning). The highest and lowest *F D*_*s*_ are represented in red and blue respectively and pointed by arrows. Here the color scale was offset by 0.5 and the midpoint was set at 0.01 for visualization. B) The *F D*_*s*_ for a dataset of 135 structures, gathered with a PSI blast seeded with a PFAM seed alignment. The structure used for FD mapping is the porcine pancreatic amylase (PDB code: 1PPI). The grey chain correspond to the non homologous section of the 1PPI with respect to the alignment. Both figures were rendered with VMD v1.91 [Humphrey et al., 1996].

Here we can see which residues contribute the most and the least to the overall shape, as well as their relative position.

As expected for the real dataset of homologs (Figure 2B), the most variable (and therefore with positive *F D*_*s*_ values) residues are in located in loops. The residue Arg10 was the most variable, and is also located at the beginning of the chain after the signal peptide; therefore its high variability may be explained by this location in the primary structure. The residue with the least variation is Phe136 which is also found in a loop. The Phe136 residue is not reported to bind to ligands or be involved in catalytic activity. However, this residue is within 15 Å (in a *C_α_ − C*_*α*_ perspective) of metal and ligand binding residues and it is also highly conserved.

### NPC1 dataset: Analysis of the residue variation in the context of ligand binding

The Niemann-Pick, Type C-1 protein (NPC1) binds cholesterol and oxysterols [Infante et al., 2008] and has an important role in the metabolism of cholesterol and other lipids. Defects in NPC1 cause malfunction of the cholesterol, sphingolipids, phospholipids, and glycolipids pathways. The protein contains 1278 residues, with 13 membrane helices and three large loops that project to the lumen of lysosomes [Infante et al., 2008]. The first luminal domain is the N-terminal domain, which comprises approximately 240 amino acids. This is a lumen domain, and therefore not in the trans-membrane region of the protein.

To check the overall contribution of each of the residues to the deformations during the simulation, equation 6 was applied and results were mapped on the original protein structure. Figure 3 shows that once the cholesterol is bound to the NPC1 (Figure 3A), most of the higher *F D* residues are not contributing to the deformation. It seems that most residues’ movement is been held in place by interactions with the ligand.

**Figure 3:**
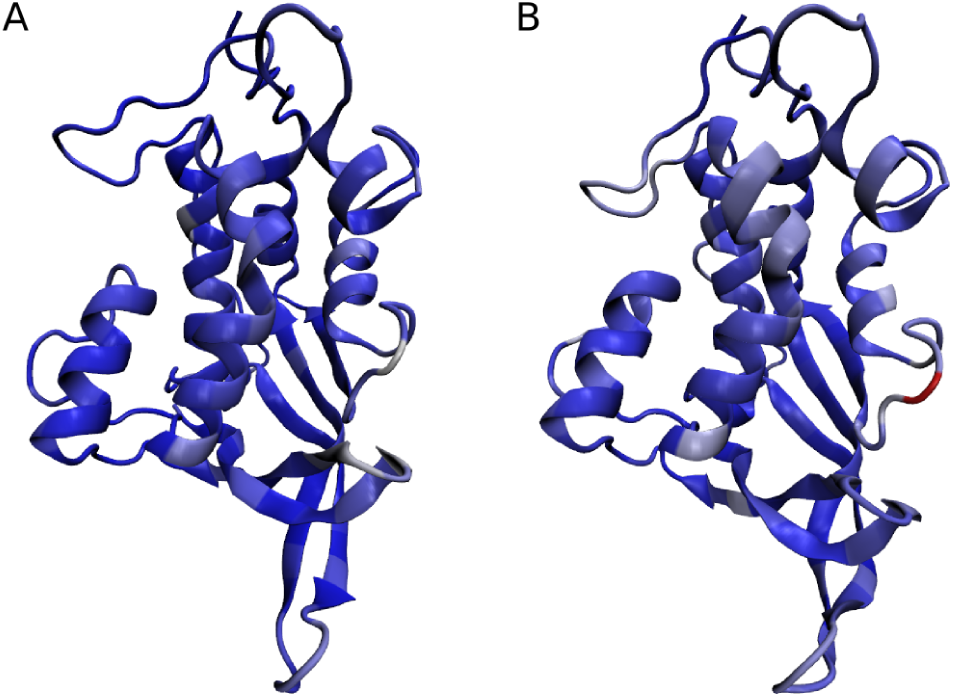
FDM analysis of the NPC1 N-terminal domain (pdbcode 3GKH) with 3A and without 3B ligand. The color represents the *F D*, red being a higher score and blue the lowest. For comparison, the scale was set from the minimum FD value in both, to the maximum between the two. In this particular case, lower *F D* values (therefore the least variables) dominate the scale and the most influential is shown in red.

The opposite behaviour can be seen in Figure 3B. When cholesterol is not bound to it, the residues in charge of the cholesterol intake/outtake are more movable. Therefore, these residues are responsible for most if the deformation. This makes sense if the binding pocket must be flexible enough to open and close upon binding with the ligand.

### GM-like analysis of protein data: An example from the multidimensional scaling

#### *α*-Amylase homologs: Geometry, function and structural similarity

After aligning the structures and applying the methods exposed in section Material and methods, a principal coordinate analysis (PCoA) was performed to the resulting landmark data (Figure 4). Analyzing the geometry of the protein structures using a PCoA can give us insight into the relationships of such shapes. This procedure test for differences in the structures being compared, and will show patterns of clustering based on their geometric similarity which in turn might be highly correlated with the functional similarity [Wright and Dyson, 1999].

**Figure 4:**
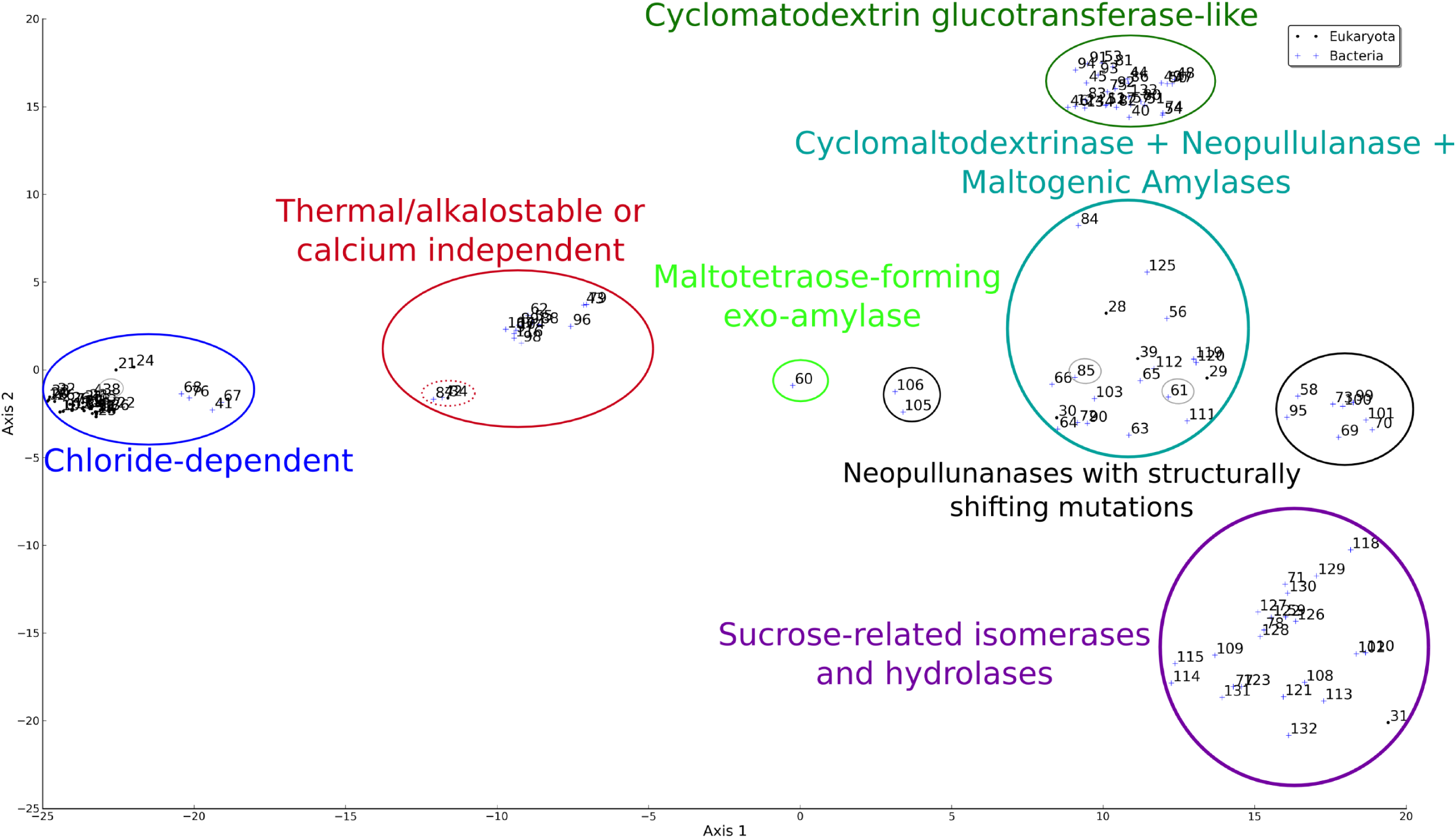
Principal Co-ordinates Analysis (PCoA) of 135 protein crystals of the *α*-Amylase. The circled groups show clusters of structural similarity. The PCoA was performed in R [R, 2011] and modifid with a python script using Matplotlib [Hunter, 2007] library.

The PCoA of the multiple structure alignment (Figure 4), showed seven distinct and tightly clustered groups:

> **Chloride-dependent** *α***-Amylases**
>
> The first group corresponds to the Cloride-dependent *α*-Amylases (with amylase function or EC # 3.2.1.1). The similarity among these *α-*amylases is not a new observation. D’Amico et al. [2000] identified potential chloride-dependent amylases, based on the chloride allosteric activation positives: A) PPA or porcine (*Sus scrofa*) pancreatic *α-*amylase; B) HPA or human (*Homo sapiens*) pancreatic *α*-amylase; C) TMA or *Tenebrio molitor* (mealworm) *α*-amylase; and D) AHA or *Pseudoalteromonas haloplanktis* (before classified as *Alteromonas*) *α-*amylase. They showed that the side chains of residues Arg195, Asn298 and Arg/Lys337 (PPA numbering), are related to chloride ion binding capabilities [Da Lage et al., 2004].
>
> **Thermal/alkalostable or calcium independent** *α***-Amylases**
>
> The next tightly defined group in Figure 4 are structures that show higher stability in extreme PH and/or thermal conditions or are calcium independent. As shown in Figure 4 there is a subgroup of mutants with higher structural shift from the main group. In this sub-cluster three thermo-stable *α*-amylases (EC # 3.2.1.1) mutants from the genus *Bacillus* can be found. In two of the three cases (3DC0, Rahimzadeh et al., 2012; 1BF2, Fujimoto et al., 1998) the directed mutagenesis was performed to increase thermal stability. In the case of 1UA7, it is a mutant of the catalytic site that is not supposed to change stability or function with respect to the wild type [Kagawa et al., 2003]. However, this structure was modeled using 1BF2, and the clustering observed in its structure suggests a higher performance or thermal-stability than other non-chloride binding bacterial amylases. The rest of the group includes *α*-amylases that exhibit higher thermal/alkaline stability or enhanced efficiency with respect to other amylases of similar function (*α*-1,4-glucan-4-glucanohydrolase, EC # 3.2.1.1) [Shimi et al., 2008]. Most of these structures were created by directed mutagenesis to enhance their industrial applicability by either increasing their thermal or alkaline resistance [Hwang et al., 1997, Machius et al., 1998, Brzozowski et al., 2000, Machius et al., 2003, Lyhne-Iversen et al., 2006, Shirai et al., 2007, Shimi et al., 2008, Alikhajeh et al., 2010] or to make them calcium independent [Prakash and Jaiswal, 2010]. There is also a structure with a different enzymatic classification, the maltohexaosidase from *Bacillus licheniformis* (1WP6; glucan 1,4-*α*-maltohexaosidase or EC # 3.2.1.98). Despite carrying a slightly different reaction, its native state exhibit higher alkaline stability than other native amylases [Kanai et al., 2004a].
>
> **Cyclomaltodextrinase-like** *α***-Amylases**
>
> The Cyclomaltodextrinase + Neopullulanase + Maltogenic Amylases group (Figure 4) includes enzymes classified in seemingly different functional groups (Cyclomaltodextrinases EC # 3.2.1.54; maltogenic amylases, EC # 3.2.1.133; neopullulanases EC # 3.2.1.135) that can hydrolyze cyclomaltodextrins efficiently [Park et al., 2000] but not starch and pullulan as efficiently [Lee et al., 2002]. However, Lee et al. [2002] have shown that despite their different enzyme codes, there are no thoroughly documented differences in the literature about their function or structure. They proposed to unify this group under the same enzyme number and the same name (Cyclomaltodextrinases). The result, shown in Figure 4, suggests that this is the case given our clustering based on shape. It is important to mention that this Cyclomaltodextrinase group has to be distinguished from the Cyclomatodextrin glucotransferase group, since those are extracellular enzymes whereas the Cyclomaltodextrinase-like *α*-Amylases are intracellular[Lee et al., 2002].
>
> **Cyclomaltodextrinase-like** *α***-Amylases with structural shifts**
>
> The Neopullunanases with structurally shifting mutations groups is a subset of the Cyclomaltodextrinases described above. They carry the same functions (mainly Neopullunanse; EC # 3.2.1.135), but have been subjected to mutagenesis either for binding studies (i.e. 2FH8 and 2FHB; Mikami et al., 2006) or to inactivate the enzyme using site-directed mutagenesis [Ohtaki et al., 2001, Yokota et al., 2001, Ohtaki et al., 2004, Mizuno et al., 2005]. As can be seen in Figure 4, even a small number of substitution cause structural shifts that can be identified by means of a PCoA.
>
> **Cyclomatodextrin glucotransferases-like** *α***-Amylases**
>
> This group is composed entirely of bacterial (mainly from the genus *Bacillus*) *α*-Amylases that catalyze the conversion of starch to cyclodextrins (EC # 2.4.1.19) [Kanai et al., 2004b]. As can be seen in Figure 4, it is a tightly defined group markedly different from the rest. These differences can be explained by the presence of four aromatic residues that are not present in other amylases and are strongly associated with the protein function [Tonkova, 1998, Kanai et al., 2004b].
>
> **Maltotetraose-forming exo-amylase**
>
> This is singleton group, containing the structure 1GCY [Mezaki et al., 2001] from *Pseudomonas stutzeri*. It is a glucan 1,4-alpha-maltotetraohydrolase (EC # 3.2.1.60) that works hydrolyzing amylaceous polysaccharides and removing successive maltotetraose residues from the non-reducing chain ends [Fleischmann et al., 2004]. It behaves as an exo-amylase and structural differences with respect to endo-amylases were expected, to be able to remove the residues at the end of the chain instead of just breaking the 1-4 glycosidic linkages. The PCoA in Figure 4 expresses this differences by showing a distance between this structure with the “endo-amylases”.
>
> **Sucrose-related isomerases and hydrolases**
>
> This group contains the structures mainly classified as Sucrose glucosyl-mutases (or Isomaltulose synthase EC # 5.4.99.11)[Fleischmann et al., 2004]. However, it also contains three structures (namely 2ZIC, 2ZID, and 4AIE), with Glucan 1,6-alpha-glucosidase (EC # 3.2.1.70) function [Kim et al., 2005, Hondoh et al., 2008, Møller et al., 2012], from which the 2ZIC and 2ZID are have been subjected to directed mutagenesis to improve their catalytic efficiency. This group also harbors an *α*-glucosidase (2ZE0; Shirai et al., 2008) and an Oligo-1,6-glucosidase (Isomaltase; 1AXH) mutant [Yamamoto et al., 2011]. The rest of structures there are Isomaltulose synthases (EC # 5.4.99.11), including three (4GIN,4GI6, and 4H2C) misannotations (inexistent EC # 5.4.11.99 instead of EC # 5.4.99.11). Despite the somewhat disparity in function, they all are classified in the GH13 family [Cantarel et al., 2009, Svensson and Janecek, 2013], and the results shown in Figure 4 suggest a high structural similarity.

As can be seen, the principal coordinate analysis of protein structures is tightly correlated with function, and might give some insights into missannotations or potential functional discoveries. This can be useful for the classification of proteins. This clustering scheme might show an apparent correlation with phylogeny. However, this approach showed to be sensitive to structural changes. It identified even mutants from the wild type if a structural shift has occurred. This may suggest that this approach is capturing more structural similarities than only phylogenetic ones. It would be interesting to explore phylogenetic signal free variables to test such a hypothesis, but even now this approach seems to be robust to find functional/structural groups.

#### NPC1 molecular dynamic data

The NPC1 N-terminal domain dataset was explored using a PCoA (Figures 5A and 5B) to analyze the trajectories as a composite measure of overall structures (principal coordinates).

**Figure 5:**
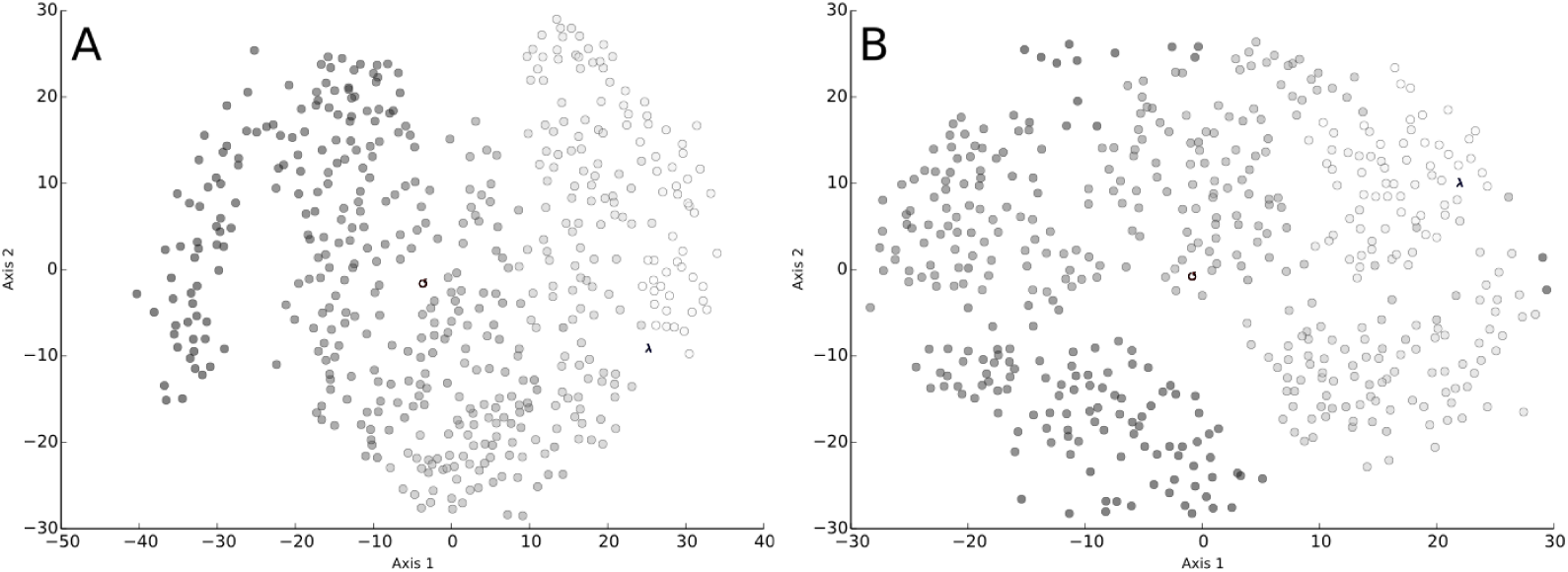
Principal coordinate analysis of 495 snapshots from 100 ns of molecular dynamic simulations of the NPC1 N-terminal domain without (5A) and with (5B) cholesterol bound to it. The gray scale is proportional to the time of the simulation getting progressively lighter as the simulation develops. The symbols ↺ and *λ* represent the starting and final points respectively

This exploratory analysis seems to indicate that a burn-in of 10 nanoseconds and a sampling every 20 ps achieves stability in both cases (Figure 5). The overall structure seems to sample the space around the initial point (Figure 5). However, the simulation with the cholesterol (5B) seems to have a narrower shift (as per first axis) than the ligand-free (5A). The shifts here are represented for a wider move in the principal coordinates space. This technique can be used to see if clusters of structures appear, and therefore a deviation from equilibria or representation of multiple states in a molecular dynamic simulation. This method in conjunction with the energy profile and a clustering scheme, can provide useful information about the dynamic properties of proteins.

## Conclusions

Here we have shown how to apply GM-like methods to the protein structure variation analysis problem. We have also shown that by using GM-like methods, it is possible to explore patterns in both evolutionary and dynamic samplings. Due to the scarcity of space, a full application of different statistical techniques could not be done. However, the application of techniques, such as linear discriminant analysis, support vector machine, random forest, multivariate analysis of variance, different types of regression, among others, are simple extensions of what we have shown here. Such techniques might help in classification of proteins (structurally or phylogenetically) as well as characterizing the variation associated with different parameters in the structural space.

## Acknowledgements

The authors thank professors Andrew Roger, Jan K. Rainey, and Edward Susko for useful insights. Also, the members of the Blouin lab for the critical review of the manuscript and to Liz Mackay for the language editing the manuscript. This study was funded by NSERC through the grant No. 120504858. This work was partially supported by The Departamento Administrativo de Ciencia y Tecnología - Colciencias (Colombia) through the CALDAS scholarship.

## Supplementary data

Table S1: PDB codes of the *α*-amylase homologues, the species from it was crystallized, and its corresponding equivalence number in the PCoA plot in Figure 4. Only the chain A, corresponding to the catalytic domain, was used.

